# CNVkit-RNA: Copy number inference from RNA-Sequencing data

**DOI:** 10.1101/408534

**Authors:** Eric Talevich, A. Hunter Shain

## Abstract

RNA-sequencing is most commonly used to measure gene expression, but it is possible to extract genotypic information from RNA-sequencing data, too. Point mutations and translocations can be detected when they occur in expressed genes, however, there are few software solutions to infer copy number information from RNA-sequencing data. This is because a gene’s expression is dictated by a number of variables, including, but not limited to, copy number variation. Here, we report new functionalities within the software package CNVkit that enable copy number inference from RNA-sequencing data. First, CNVkit removes technical variation in gene expression associated with GC-content and transcript length. Next, CNVkit assigns a weight, dictated by several variables, to each transcript with the net effect of preferentially inferring copy number from highly and stably expressed genes. We benchmarked our approach on 105 melanomas from The Cancer Genome Atlas project and observed a high degree of concordance (R = 0.739) between our estimates and those from array comparative genomic hybridization (aCGH) on the same samples. After initial configuration, the software requires few inputs, is able to process a batch of up to 100 samples in less than ten minutes, and can be used in conjunction with pre-existing features of CNVkit, including visualization tools. Overall, we present a rapid, user-friendly software solution to infer copy number information from gene expression data.

## Introduction

RNA sequencing is predominantly utilized to measure gene expression, while DNA-sequencing is utilized to measure genotypes. It is informative to have both sets of information to investigate genotype-phenotype relationships, but matched DNA- and RNA-sequencing from the same sample is expensive and often not possible to perform. In particular, droplet sequencing technologies have become popular because they enable parallel RNA-sequencing from thousands of individual cells, but the library preparation techniques for droplet sequencing are not compatible with matched DNA-sequencing^1–3^, prohibiting investigations into genotype and phenotype relationships from these experiments.

An alternate solution to this challenge is to directly assess genotypes from RNA-sequencing data. Some genotypic information, such as point mutations and translocations, can be detected in RNA-sequencing data if they occur in expressed genes. In contrast, copy number alterations are difficult to detect from gene expression data because each gene’s expression is dictated by a number of variables, including, but not limited to, its copy number. This is a significant shortcoming, particularly when studying cancer cells, because it is common for thousands of genes to have copy number aberrations that deviate from the diploid state.

There are virtually no software solutions to infer copy number from gene expression data. Regev and colleagues utilized a clever approach in which they averaged the expression of a moving window of 100 adjacent genes, reasoning that concordant up- or down-regulation of large numbers of neighboring genes would most likely be driven by copy number alterations^4^. This enabled them to differentiate stromal cells and tumor cells, which respectively have no copy number alterations or many copy number alterations, from their single-cell RNA sequencing data, exemplifying a particularly useful application for this type of analysis. Here, we have substantially improved upon their basic methodology, described below and implemented as a new module in the software package CNVkit^5^.

## Methods

### Workflow

CNVkit operates under the basic assumption that a gene’s expression is correlated with its copy number. However, other biological factors influence gene expression, and technical variables can affect the precision and accuracy of gene expression measurements. To address these challenges, CNVkit removes technical biases and infers copy number preferentially from stably expressed genes, highly expressed genes, and genes whose expression is known to be closely connected to copy number (Fig. 1).

**Figure 1.**
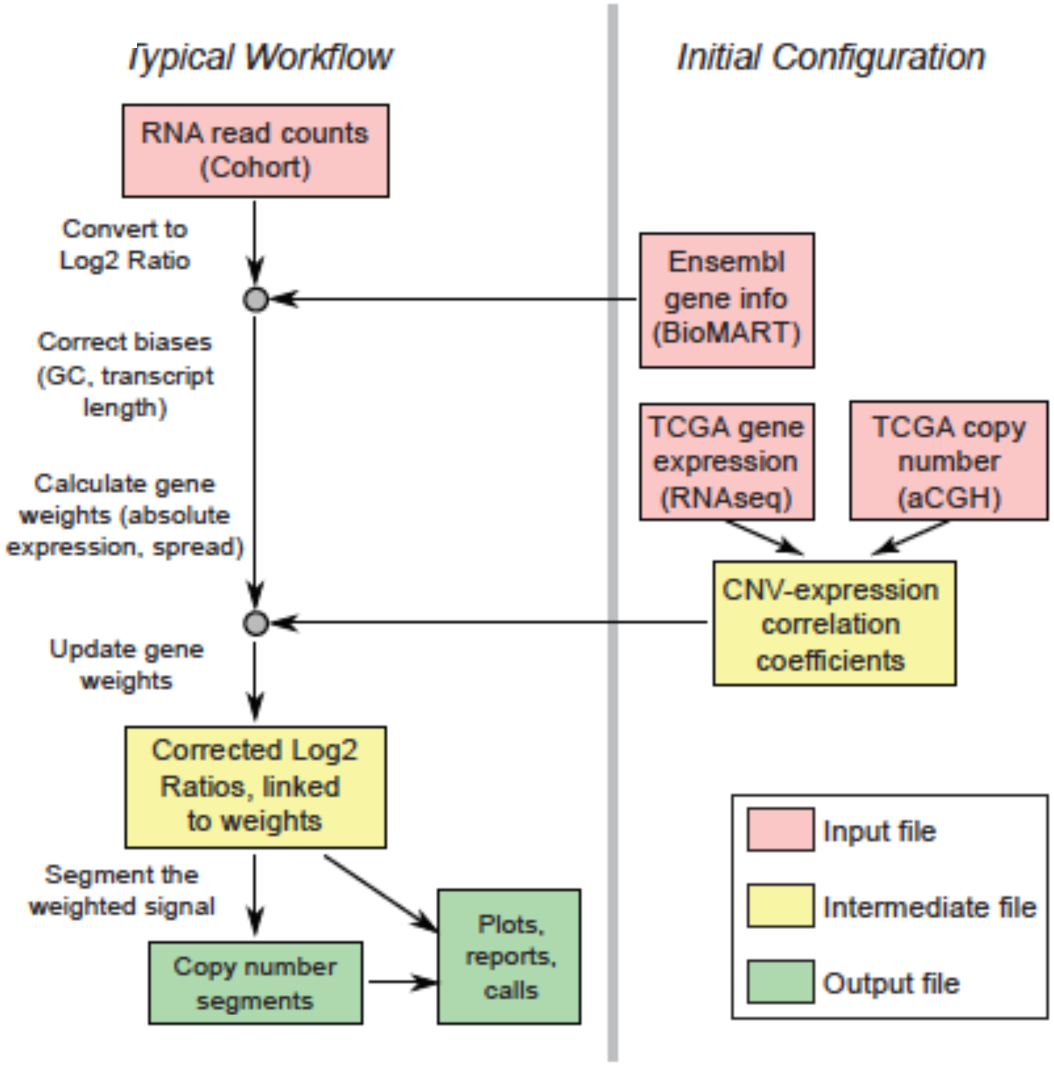
The CNVkIt wokflow to infer copy number from RNA-Sequencing data.

### Inputs and Preprocessing

CNVkit is designed to operate on a batch of samples for which gene expression data is available. We recommend batching samples from similar cell types. This is because some genes are specifically expressed in certain cell lineages, and their expression may correlate well with copy number within a lineage but not across lineages.

The basic input for CNVkit is a two-column table with gene IDs and sequencing read counts. In lieu of this table, CNVkit can accept the output of RSEM^6^, a popular software program that quantifies read counts per gene, aggregated across transcripts. CNVkit also requires a resource file that links transcript IDs to other variables necessary for copy number inference. A general-purpose resource bundle is included in the software download.

To begin, CNVkit executes several preprocessing operations. First, each gene is linked to its matching features from the resource bundle, while genes that are not expressed in the majority of samples are filtered out. Read counts are then normalized within each sample, controlling for differences in sequencing depth across the samples, and normalized by gene, controlling for differences in basal levels of expression across genes. Ultimately, this step produces a tabular file of log_2_ expression ratios for each gene.

### Account for known systematic biases in gene expression

The number of sequencing reads corresponding to a gene is primarily influenced by biological factors but can be skewed by technical factors. For instance, the efficiencies of reverse transcriptase and DNA polymerase enzymes, which are utilized during library preparation, are affected by the GC content of each transcript^7^ (Fig. 2). Further, differences in RNA quality can influence the resulting sequencing counts, particularly for longer transcripts, which are known to be less stable and degrade from the 3’ end^8^. CNVkit improves the accuracy of read counts by removing expression biases, associated with GC content or transcript length, within each sample using a rolling-median correction.

**Figure 2.**
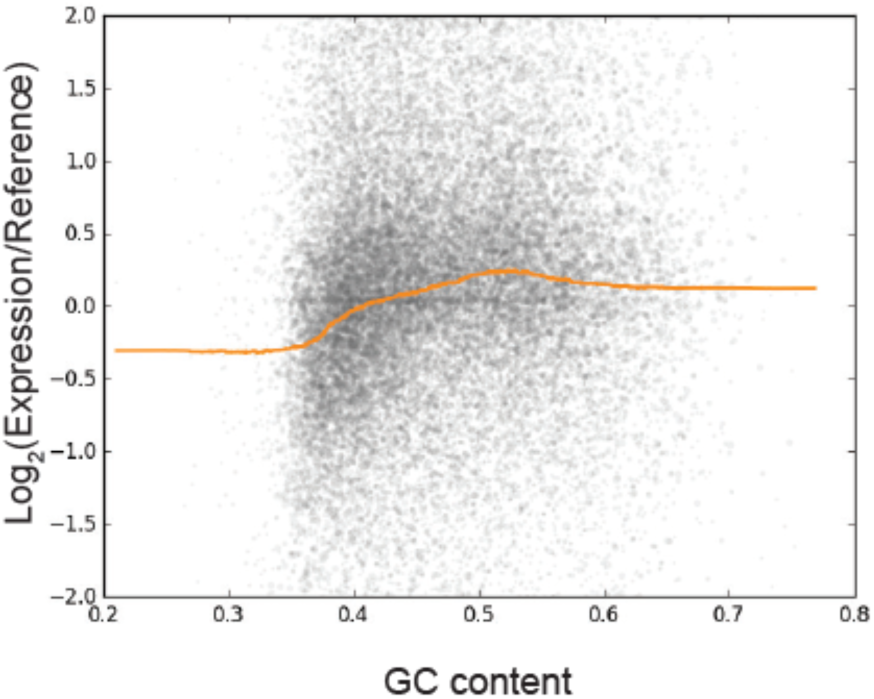
An example of expression biases introduced by GC-content. Each grey data point corresponds to a gene stratified by its expression (Log2 scale, y-axis) and GC content (x-axis). The yellow lines denotes a moving average. Note the the systematic biases in gene expression correlating with GC content. CNVkit removes these biases, which are assumed to be purely technical, with a rolling median correction. This data is from a single sample in the melanoma cancer genome atlas project.

### Weight genes by reliability of copy number signal

GC-content and transcript length introduce biases in gene expression that can be measured and thus removed. However, the specific effects on gene expression of many other variables are not possible to measure, prohibiting precise quantification of variability in gene expression driven by copy number. To address this challenge, we assign a weight to each gene in an attempt to quantify each gene’s ability to provide copy number information. Subsequent steps apply these weights in order to preferentially use the more reliable genes to infer and visualize copy number information.

Absolute gene expression is the first variable to influence the weight of each gene. Only a subset of the transcriptome is ultimately sequenced, and this random sampling can produce over- or under-estimation of gene expression measurements, exemplifying a bias that exists but for which the magnitude and direction is impossible to measure (Fig. 3). In particular, random sampling biases are exacerbated in poorly expressed transcripts. For instance, a transcript that should have 5 reads at a given sequencing depth could be under- or over-sampled, resulting in a measurement that misses by at least 20%, and possibly much more. CNVkit reduces the weight for poorly expressed transcripts to reflect the fact that their measurements are less precise than measurements from highly expressed transcripts.

**Figure 3.**
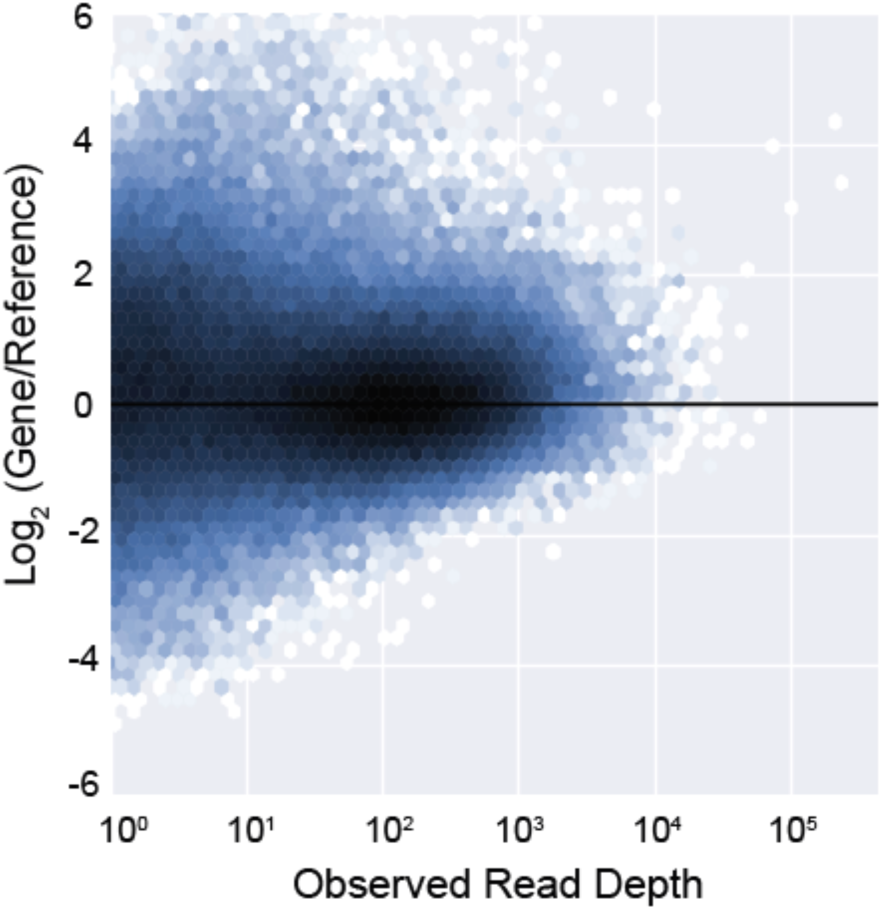
Highly expressed genes tend to have more stabe expression. Density plot stratifying all genes ¡n all samples from the melanoma TCGA project by their absolute expressions (x-axis) and normalized expressions (y-axis). Note that the range of normalized expression decreases for more highly expressed genes.

The variablity of gene expression is the second variable to influence the weight of each gene. Many genes are expressed over a dynamic range, but the range of copy number variation tends to be much smaller. Therefore, for each gene CNVkit calculates the standard deviation of normalized expression ratios across the batch of samples being tested and adjusts each gene’s weight to favor more stably expressed genes. This adjustment assumes that stably expressed transcripts are more reliable indicators of copy number.

Finally, CNVkit leverages pre-existing data to identify genes whose expression is known to correlate with copy number. We downloaded matched expression and copy number data from The Cancer Genome Atlas project and calculated correlation coefficients between expression and copy number for each gene. The expression of many housekeeping genes correlates well with their copy number. For instance, *TBP*, or TATA-Box Binding Protein, encodes a component of the RNA Polymerase II transcription complex, and its expression is driven almost entirely by copy number (Fig. 4). In contrast, the expression of other genes, including many transcription factors such as *SOX17* (Fig. 4), show poor correlation, as their expression is presumably modulated by other signals. We include our pre-calculated correlation coefficients as part of the CNVkit software distribution. For users who wish to derive their own coefficients from other datasets, CNVkit also includes a script to calculate correlation coefficients from any dataset with matching gene expression and copy number data.

**Figure 4.**
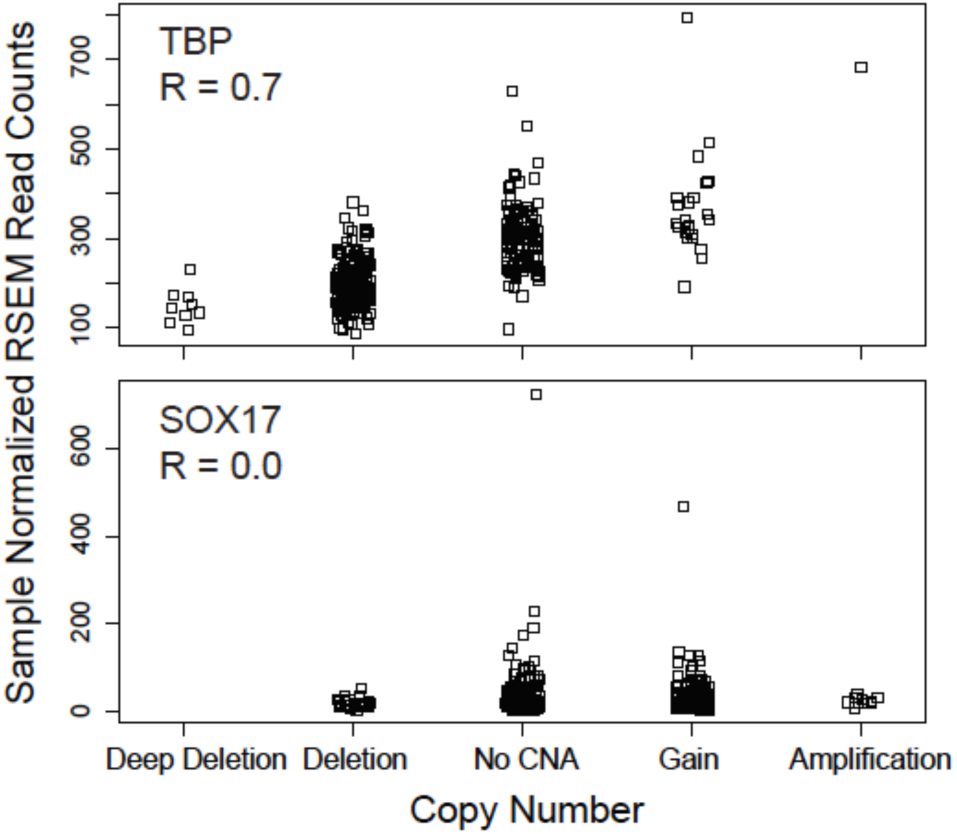
The proportion of gene expression dicated by copy number varies across genes. TBP, or TATA-binding protein, is a housekeeping gene whDse gene expression is dicated primarily by copy number, as shown across melano ma cancer genome atlas samples. In contrast, SOX17 is a transcription factor whose expression is primarily dictated by other factors.

The net effect of these weight adjustments is to favor highlyand stably-expressed housekeeping genes when inferring copy number information from gene expression measurements.

### Segmentation

Copy number is ultimately inferred from the gene expression data by using a segmentation algorithm that can incorporate the weights described above. CNVkit utilizes circular binary segmentation (CBS)^9^ by default, though other segmentation algorithms that incorporate weights could instead be utilized.

## Benchmarking

We benchmarked our data on 105 melanomas from the Cancer Genome Atlas project. These samples were profiled by RNASequencing and by array comparative genomic hybridization (aCGH), the gold standard by which we compared our copy number inferences. All supporting files and scripts for these analyses are available online at https://github.com/etal/cnvkit-examples. These 105 samples were not included in the correlation value calculation described above.

A comparison of copy number segments inferred by CNVkit and by aCGH are shown in figure 5 for an individual sample, illustrating the strengths and limitations of our approach. CNVkit was capable of detecting broad copy number alterations; however, focal alterations, such as a homozygous deletion of *CDKN2A* on chromosome 9, were sometimes missed. This is not surprising because the Affymetrix SNP 6.0 array, utilized by The Cancer Genome Atlas project, has nearly 2 million probes, whereas there are only ∼24,000 genes, many of which are not expressed. Therefore, the resolution to detect copy number alterations from RNA-sequencing is limited when compared to high density arrays.

**Figure 5.**
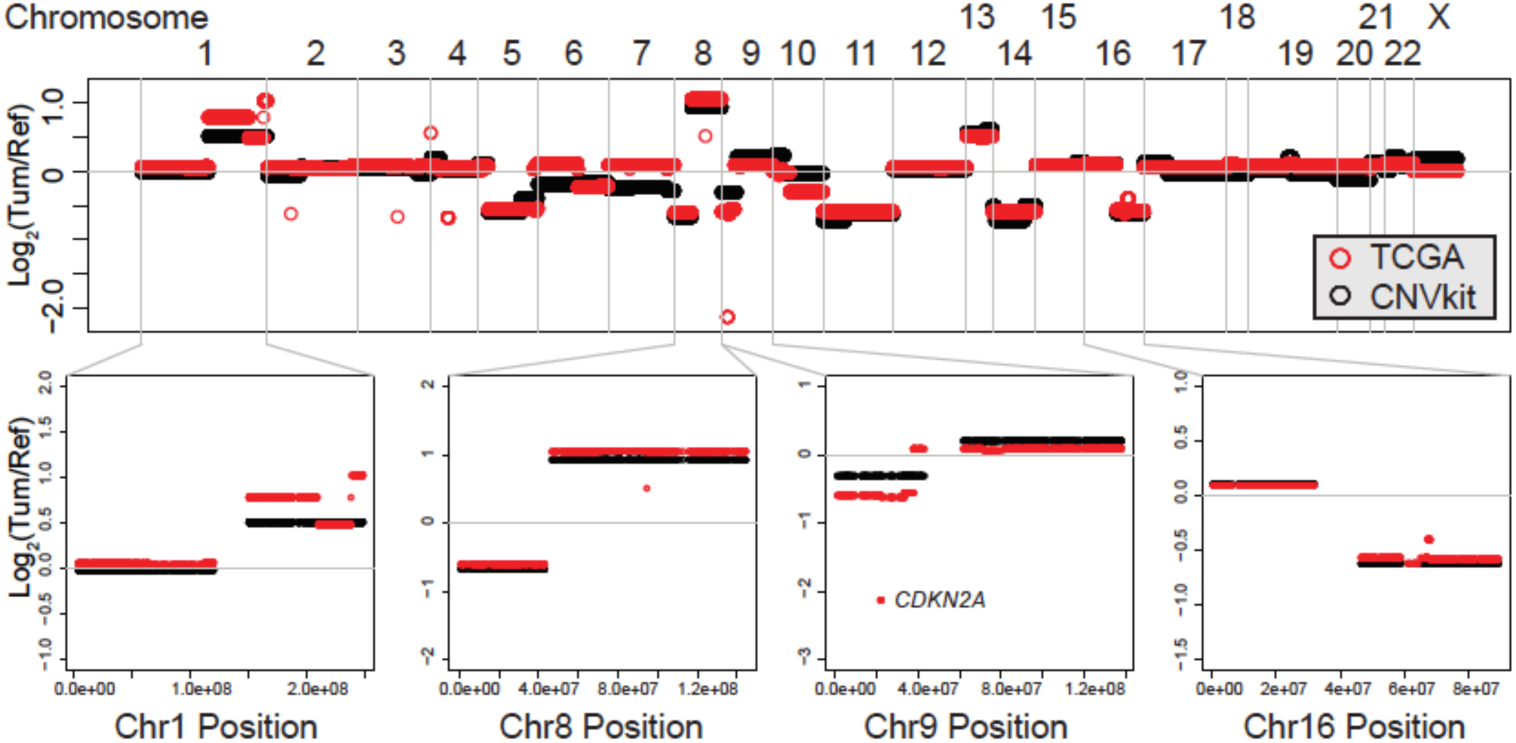
CNVkit copy number inference from RNA-Sequencing counts is concordant with aCGH copy number inference. Copy number estimates for all genes in a single sample from the melanoma cancer genome atlas project Each individual circle corresponds to a gene, plotted by its position along the genome and inferred copy number TCGA calls (red) stem from segmented aCGH data (Afíymetrix SNP 6 0) and CNVkit calls (black) were inferred from RNA-sequencing counts as deScribed. Several chromosomes are shown in zoomed insets to illustrate the perîorrnance of CNVkIt.

Next, we investigated the improvements associated with bias corrections, gene weighting, and segmentation (Fig. 6). To do this, we first inferred copy number by averaging the expression of adjacent genes, as described by Tirosh *et. al.*^4^, and compared these estimates to copy number measurements made by aCGH from the same samples (‘Smoothing’ method in Fig. 6). We sequentially introduced optimizations to measure the improvement associated with each. Bias correction, gene weighting, and segmentation resulted in copy number estimates that were closer to those obtained by comparative genomic hybridization (Fig. 6). These improvements held true for large-(>50 Megabase, Mb), medium-(5-50Mb), and small-(<5Mb) scale copy number alterations.

**Figure 6.**
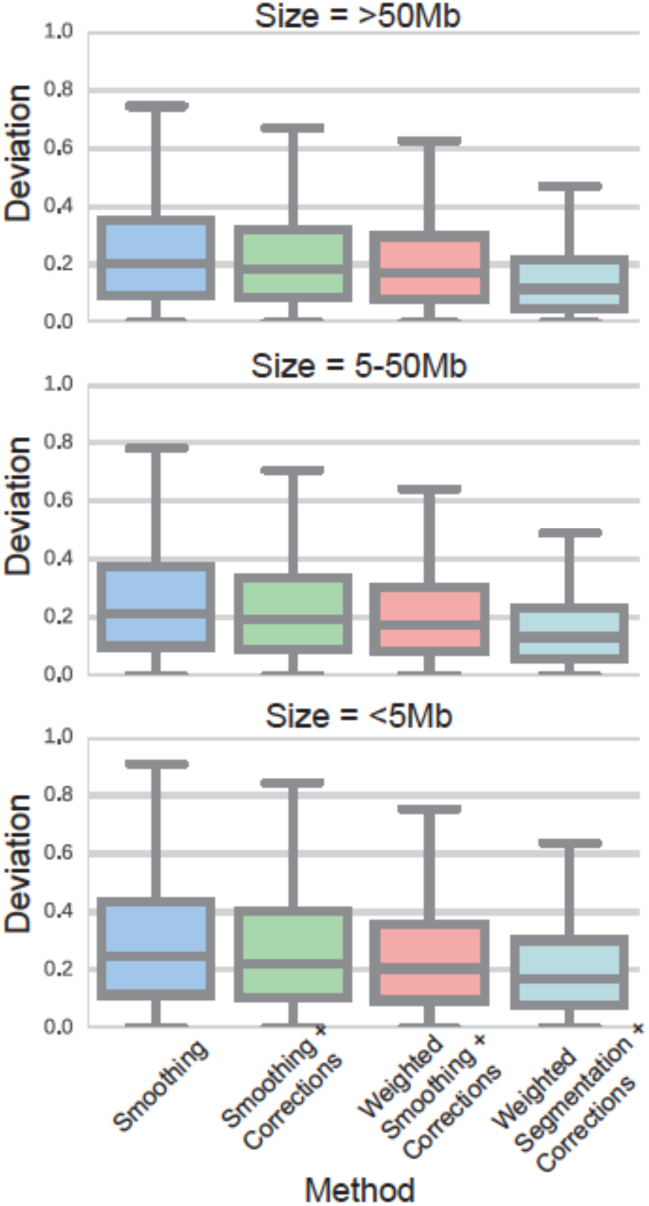
Bias correction, gene weighting. and segmen tation improve copy number inference from RNA-sequenc ing data. We inferred copy number information from RNA-sequencinç data for all genes from 105 melanomas in the cancer genome atlas project and compared our estimates to copy number measurements by array comparative genomic hybridization (aCGH, Affymetrix SNP6.0 platform) from the same tumors. The y-axis represents the relative distance between our estimates and those made by aCGH (log2 fold change). We made estimates under four conditions: simple averaging of 100 adjacent genes (as described by Tirosh et. al. 2016), simple averaging after correcting for GC content and transcript length, weighted averaging of corrected data, and weighted segmention of corrected data. Improvements were observed by each of these modifications for small (<5 Megabase, Mb), medi um(5-5OMb), and Iarge(>5OMb) scale copy number alterations.

## Output

From RNA-sequencing counts, CNVkit produces probe-level and segment level copy number estimates – respectively output as “.cnr” and “.cns” files. These file formats mimic the output of CNVkit’s DNAsequencing workflow, and therefore they are compatible with the many pre-existing visualization tools built into the software suite. For instance, CNVkit can produce scatterplots of copy number alterations, as shown (Fig. 7). In these scatterplots, the size of each datapoint (gene) is proportional to the gene’s weight, thus indicating which genes primarily drove segmentation. In the example scatterplot shown in figure 7, note that the large datapoints, which have higher weights, tend to reside closer to copy number segments. A full listing of visualization tools are described here (https://cnvkit.readthedocs.io/en/stable/). A tutorial on how to run the software is available here (https://youtu.be/JGOIXYoLG6w).

**Figure 7.**
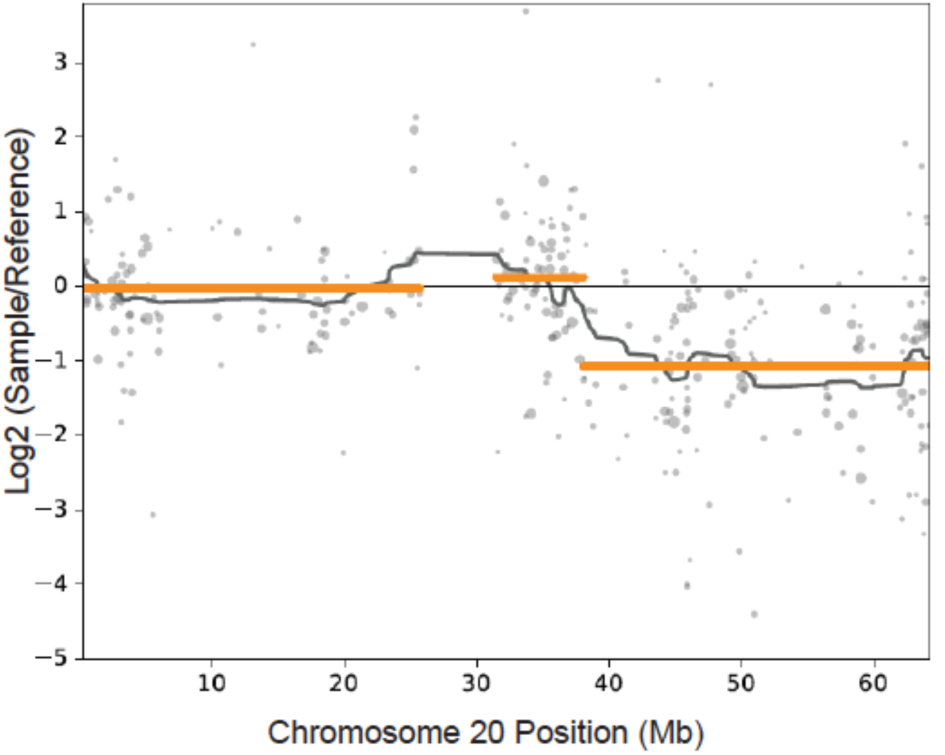
Visualization of a copy number loss on chromosomal arm 20q. CNVkit is capable of outputting data in many forms and includes built-in visualization tools Shown here is a scatterplot of a deletion on chromosomal arm 20q Each grey data point corresponds to the expression level of an indivdual gene (after corrections). The size of each data point is proportional to the gene’s weight, or reliability. Genes with higher weight primarily drove clustering and therefore reside closer to the copy number segments (gold lines).

## Discussion

Here, we present a software solution to infer copy number information from RNA-sequencing data as a new module to the CNVkit software package. CNVkit was originally designed to infer copy number information from DNA-sequencing data. For DNA-sequencing data, read depth is dictated by copy number variation as well as technical factors, such as GC content; therefore, to infer copy number information from DNA-sequencing data, CNVkit removes technical variation from read depth and infers copy number information from the remaining variability in read depth. Many of the same technical factors influence read depth within RNAsequencing data, and we use the existing framework of CNVkit to remove these sources of variability in RNAsequencing data, too. However, RNA-sequencing data poses an additional challenge because the remaining variability in gene expression is dictated by a number of biological variables including, but not limited to, copy number variation.

To address this challenge, CNVkit assigns a weight to each gene that attempts to capture each gene’s ability to provide copy number information. To calculate this weight, CNVkit leverages information from the RNAsequencing dataset itself, including the absolute expression of each gene and variability in the expression of each gene across the batch of samples being studied. CNVkit assumes that the expression of highly- and stably-expressed genes is more likely to reflect the number of copies of those genes. CNVkit also leverages large-scale, independent datasets of matching gene expression and copy number data to calculate the weight of each gene. Altogether, CNVkit preferentially weighs stably- and highly-expressed genes whose expression is known to concord with copy number in order to segment gene expression data.

There are numerous applications for this software. For instance, thousands of gene-expression profiling studies already exist, and this software can be used to go back and extrapolate copy number information from the samples in those studies. As a recent example^10^, we used this software to identify deletions of chromosome 3, harboring the *BAP1* gene, in melanocytic neoplasms for which we hypothesized that *BAP1* was deleted but only RNA-sequencing data existed.

Moving forward, this software will be useful to extract genotypic information, at no additional cost, from RNAsequencing studies when it is not feasible to also collect DNA-sequencing data. In particular, this will be useful for single-cell RNA-sequencing studies because it is challenging to extract DNA and RNA from the same cell. Overall, CNVkit makes it possible to collect genotype- and phenotype-information from a single assay.

In closing, the import-rna command of CNVkit, described here, is part of a larger software suite that is actively maintained. This software is user-friendly and can be executed to process hundreds of samples in a manner of minutes. The output of this command pairs with the downstream visualization tools that are part of the original CNVkit distribution. CNVkit now provides production-quality software to infer copy number information from both DNA- and RNA-sequencing data.

## Supplemental Methods

### Program inputs

A tutorial that walks through the basic implementation of the import-rna command is available here, https://youtu.be/JGOIXYoLG6w, and described in greater detail below.

The ‘import-rna’ command takes a batch of samples’ per-gene RNA sequencing read counts as the primary input. At minimum, for each sample, this consists of a 2-column tabular file listing each Ensembl gene name and the number of sequencing reads counted for that gene. Alternatively, the gene-level RSEM output file may be used. In the latter case, the estimated (weighted-average) transcript length for each gene is also extracted to be used in the bias-correction step.

The command also requires a separate table of metadata for each gene: Ensembl gene ID, G+C percentage, chromosome name, start and end genomic coordinates, HGNC gene name, NCBI Entrez gene ID, transcript length, and transcript support level. This table can be retrieved from Ensembl BioMart. For convenience, a table for the human genome build hg38 is included in the CNVkit source distribution.

### Calculation of copy-number-to-expression correlation coefficients

An optional but strongly recommended input is a table of each gene’s correlation of expression and copy number as observed in a separate study. Summary tables of this information are publicly available for TCGA studies, where these values were measured by RNA sequencing and high-density array CGH.

To calculate per-gene correlation coefficients from TCGA datasets (or equivalent), CNVkit provides a script ‘cnv_expression_correlate.py’. For convenience, the correlation table for The Cancer Genome Atlas melanoma project has been precomputed and is included in the CNVkit source distribution.

### Expression ratio normalization

After ingesting all sample’s per-gene read counts, the ‘import-rna’ command identifies and discards genes that are not consistently expressed with a count of at least 1 in a majority of the given samples. Genes that do not appear in the Ensembl gene metadata table are also discarded.

To account for variations in sequencing depth between samples, and relative differences in expression between genes, the per-gene read counts are centered by sample and by gene, iteratively. Since most genes will have low expression in a given sample, we follow TCGA guideline to improve the stability of sample-wise centering by taking the third quantile (75%) of read counts to divide the sample’s read counts. For gene-wise centering, we divide read depths by the median across all samples for each gene. After centering, the normalized expression ratios are converted to log2 scale. The result is a log2-scaled expression ratio for each gene in each sample, comparable across samples.

In some studies, there is a risk that some copy number alterations are present in a large fraction or even a majority of samples. To guard against these copy number alterations influencing the centering routine described above, the ‘import-rna’ command can also accept a list of “normal” or control samples, preferably from the same batch as the test samples, but that are not expected to contain recurrent CNAs. If given, the control samples are included in the gene-wise and sample-wise centering routine along with the test samples. Then, after centering is complete, the test samples’ expression ratios are divided by the median of the control samples’, nullifying any shift introduced by recurrent CNAs in the test samples. (An alternate approach of dividing out the the upper and lower quartiles of the normal samples, rather than the median, as described by Tirosh et al.^4^, is implemented but not enabled by default.)

### Calculation of per-gene weights

The ‘import-rna’ command assigns each gene a weight between 0 and 1, indicating its reliability for copy number estimation. The weights are calculated from several sources in the program inputs.

Before normalizing expression ratios, CNVkit calculates each gene’s median read count across samples. Considering that many genes have low expression, as in the centering routine described above, the upper quartile (75%) of per-gene median read counts is calculated as a “well-expressed” baseline; genes with median read counts below this value are therefore down-weighted quadratically:

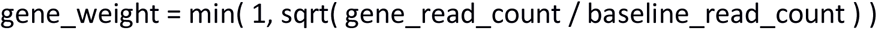

After gene-wise and sample-wise normalization, CNVkit calculates the standard deviation of log2 expression ratios across samples for each gene, resulting in a second set of per-gene weights.

If correlation coefficients are provided as input to ‘import-rna’, CNVkit uses these as a third set of weights. Pearson r, Spearman rho, and Kendall tau correlation coefficients are all accepted in the input per-gene correlation table. If multiple sets of correlation coefficients are provided, the arithmetic mean of all values given for each gene will be used.

Finally, the two or three sets of per-gene weights (depending on whether correlations were provided) are consolidated into a single column of per-gene weights by taking geometric mean across weight-sets within each gene. This approach takes a mildly pessimistic view of gene reliability in scenarios for which the weight sets disagree: the geometric mean will be lower than the arithmetic mean, but higher than the minimum of the three values.

### Correction of systematic biases

The automated correction of systematic biases implemented in CNVkit for DNA samples is applied similarly to RNA samples to correct for biases due to GC content and transcript length. To identify the relationshi pbetween each factor (GC or transcript length), per-gene log2 expression ratios are sorted by that factor’s numeric values, and a rolling-median curve is fit, similar to a LOWESS but robust to outliers. The fitted curve’s values are then subtracted from the corresponding genes’ log2 expression ratio values to remove the systematic bias.

### Segmentation

The per-gene tabular files produced by the ‘import-rna’ command can be used similarly to the per-bin or per-probe copy ratio files used by CNVkit to represent DNA sequencing and array CGH data. CNVkit’s ‘segment’ command can therefore be used to group the normalized gene expression ratios into piecewise-constant copy number regions.

Circular binary segmentation (CBS), the default segmentation method in CNVkit, performs well for most use cases involving RNA samples. Fused Lasso and HMM-based segmentation methods are also supported, but were not evaluated in this manuscript.

### Downstream analysis tools

CNVkit provides the same sub-commands for downstream analysis, visualization, and summary statistics for RNA samples as for DNA. In particular, CNVkit’s ‘scatter’ sub-command was used to create Figure 7 in this manuscript using the per-gene log2 expression ratios and the CBS segmentation of the same data file.

